# Delayed viral clearance and altered inflammatory responses resulted in increased severity of SARS-CoV-2 infection in aged mice

**DOI:** 10.1101/2024.10.03.616431

**Authors:** Émile Lacasse, Isabelle Dubuc, Leslie Gudimard, Ana Claudia Dos Santos Pereira Andrade, Annie Gravel, Karine Greffard, Alexandre Chamberland, Camille Oger, Jean-Marie Galano, Thierry Durand, Éric Philipe, Marie-Renée Blanchet, Jean-François Bilodeau, Louis Flamand

## Abstract

Since the onset of the COVID-19 pandemic, advanced age has emerged as a major predictor of disease severity. Epidemiological investigations consistently demonstrate an overrepresentation of the elderly in COVID-19 hospitalizations and fatalities. Despite this, a comprehensive understanding of the molecular mechanisms explaining how old age constitutes a critical risk factor remains elusive. To unravel this, we designed an animal study, juxtaposing the course of COVID-19 in young adults (2 months) and geriatric (15-22 months) mice. Both groups of K18(hACE2) mice were intranasally exposed to 500 TCID_50_ of the SARS-CoV-2 Delta variant with a variety of outcomes assessed on days 3, 5, and 7 post-infections (DPI). Analyses included pulmonary cytokines, RNA, viral loads, lipidomic profiles, and histological assessments, with a concurrent evaluation of the percentage of mice reaching humane endpoints. The findings unveiled notable distinctions between the two groups, with aged mice exhibiting impaired viral clearance at 7 DPI, correlating with diminished survival rates together with an absence of weight loss recovery at 6-7 DPI. Additionally, elderly-infected mice exhibited a deficient Th1 response characterized by diminished productions of IFNg, CCL2, CCL3, and CXCL9 relative to younger mice. Furthermore, mass-spectrometry analysis of the lung lipidome indicated altered expression of several lipids with immunomodulatory and pro-resolution effects in aged mice such as Resolvin, HOTrEs, and NeuroP, but also DiHOMEs-related ARDS. Collectively, disease severity implies a dysregulation of the antiviral response in elderly-infected mice relative to younger mice, resulting in compromised viral clearance and a more unfavorable prognosis. This underscores the potential efficacy of immunomodulatory treatments for elderly subjects experiencing symptoms of severe COVID-19.

**Author summary:** In this study, we investigated why older age is linked to more severe COVID-19 outcomes by comparing the progression of the disease in young (2 months) and elderly (15-22 months) K18(hACE2) mice infected with the SARS-CoV-2 Delta variant. After exposing both groups to the virus, we assessed various factors such as viral loads, immune responses, and lipid profiles in the lungs at different time points. Our findings revealed that elderly mice struggled to clear the virus by day 7 post-infection, leading to higher mortality rates and poorer recovery compared to younger mice. Aged mice showed weaker immune responses, with reduced production of key antiviral proteins like IFNg and certain chemokines. Lipid analysis also highlighted differences in molecules involved in immune regulation and lung protection, such as decreased levels of pro-resolving lipids and increased lipids associated with lung injury. These results suggest that older mice have a compromised antiviral defense, which could inform new therapeutic approaches for elderly patients with severe COVID-19.

## Introduction

In May 2023, the World Health Organization (WHO) declared that COVID-19 is no longer a Public Health Emergency of International Concern (PHEIC) as the disease switched from pandemic to endemic. However, COVID-19 is responsible for approximately 2000-14000 monthly deaths in 2023 according to WHO despite the broad vaccination campaigns. The disease therefore remains an important medical concern.

The etiologic agent causing COVID-19 is SARS-CoV-2 belonging to the Betacoronaviruses genus [1, 2]. The SARS-CoV-2 use the Angiotensin-Converting Enzyme 2 (ACE2) as a cellular receptor [3]. The presence of transmembrane protease serine protease-2 (TMPRSS-2), cathepsin L (CTSL), and Furin proteases that cleave the virus’ spike protein have been shown to be important for the entry in the host cell [3-5]. Induction of cellular death via the proptosis pathway by the virus leads to the release of damage-associated molecular patterns (DAMPs) and pathogen-associated molecular patterns (PAMPs)[6, 7]. Recognition of those molecules via pattern recognition receptors (PRRs) such as RIG-I-like receptors, Toll-like receptors, NOD-like receptors, or AIM2-like receptors leads to the production of several pro-inflammatory and antiviral mediators [8-14].

The activation of inflammatory and antiviral pathways through PRRs leads to the production of several cytokines, chemokines, and interferons as well as bioactive lipids [15-17]. Those mediators contribute to the early leucocyte recruitment that implicates monocyte/macrophage, dendritic cells, and Natural Killer (NK) cells which fuel the inflammatory environment and contribute to viral management [18, 19]. This environment could lead to the build-up of an efficient T cell response which will mediate the viral clearance and the inflammation resolution. On the other hand, a T cell lymphopenia and inability to control the inflammatory responses despite the elimination of the virus would lead to the development of severe COVID-19 [20].

Rapidly after the COVID-19 pandemic declaration by the World Health Organization, it became clear that not everyone was equally susceptible to the disease. The main risk factor that was reported was age. Several epidemiologic studies revealed that the COVID-19 fatality rates increase with age going from 1-3 % for patients below 50 years to 10-15% for 70-79 years and 15-25% for the above 80 years [21, 22]. Patients over 60-65 years of age represent about 70-80% of COVID-19 hospitalizations and deaths. A need for a more precise understanding of the disease course in the elderly population is required.

In the current study, we used the K18-hACE2 mouse model to compare the susceptibility of young adults and an elderly mouse cohort to pathogenic SARS-CoV-2 infection. Our results show that aged mice experienced were more severely affected by COVID-19 than younger mice. Old mice experienced a prolonged viral persistence with deep profiling of the mouse lung lipidome revealed a dysregulation in the timing and balance of the pro-inflammatory and immune-modulatory signals. In line with this finding, older mice exhibited impaired Th1-like chemokines and type II IFN response. In summary, our results suggest that the age-related severe COVID-19 was caused by a weaker early immune response that led to delayed viral clearance and increased lung inflammation and morbidity.

## Materials and methods

### Cell culture and virus

Vero cells were purchased from the American Type Culture Collection (Manassas, VA, USA). This cell line was passaged twice a week and cultured in Medium 199 (Multicell Wisent Inc.) supplemented with 10 % fetal bovine serum (FBS) (Corning Cellgro, Manassas, VA, USA), 10mM HEPES pH 7.2, 1mM of sodium pyruvate (Corning Cellgro) and 5μg/mL of Plasmocin® (Invivogen, San Diego, CA, USA) to prevent mycoplasma contamination. Cells were grown at 37°C with 5% CO2. The SARS-CoV-2 Delta strain was obtained from the BC Centre for Disease Control ([BCCDC] Vancouver, BC, Canada). SARS-CoV-2 was propagated on Vero cells. The infectious titer of viral preparations was 1.04×10^6^ Tissue Culture Infectious Dose 50_/mL_ (TCID_50_/mL). Viral titration was performed as described [23]. Experiments involving infectious SARS-CoV-2 viruses were performed under biosafety level 3 conditions.

### Mice

B6.Cg-Tg(K18-hACE2)2Prlmn/J mice were purchased from the Jackson Laboratories (Bar Harbor, ME, USA) and then bred in a colony at the Centre de Recherche du Centre Hospitalier Universitaire de Québec-Université Laval animal facility. Male and female mice with a mean age of seven weeks or eighty weeks were infected with 25μL of M-199 media containing 5×10^2^ (TCID_50_) of SARS-CoV-2 or 25μL of M-199 media for mock-infected mice. Mouse weight was recorded every day until euthanasia. Mice were sacrificed on days 3, 5, and 7 post-infections. A small lobe (approximatively 4mg) of the right lung was used for RNA extraction, the whole left lung for histological analysis and remaining right lung lobes (approximatively 200mg) for tissue homogenisation. Briefly, lung tissue was homoginized using the Bead Ruptor Bead Mill homogenizer (Omni, Kennesaw, GA, USA) then centrifuge at 3000xg for 20min at 4°C. Homogenate supernatant was use for cytokine analysis, infectious titer (TCID50_/mL_) analysis and lipidomic analysis.

### Reverse transcription digital PCR and quantitative real-time PCR analysis

RNA from mouse lungs was extracted using the Bead Mill Tissue RNA Purification Kit and the Bead Ruptor Bead Mill homogenizer (Omni). Following extraction, residual DNA was removed by treating the samples with DNAse I (Roche, Mississauga, ON, Canada). RNA was reverse transcribed to cDNA using SuperScript™ IV VILO™ master mix (ThermoFisher Scientific, Waltham, MA, USA)). SARS-CoV-2 viral RNA loads were determined using Droplet Digital PCR (ddPCR) supermix for probes without dUTP (Bio-Rad Laboratories Ltd, Hercules, CA, USA) and the QX200 Droplet Digital PCR System Workflow (Bio-Rad Laboratories Ltd). Quantitative real-time PCRs (qPCR) were performed on cDNA produced for RT-ddPCR analysis using the SsoAdvanced Universal SYBR Green Supermix (Bio-Rad Laboratories Ltd) on the Rotor-Gene Q 5plex (Qiagen). Primers and probes are listed in the supplementary table 1.

### Histological Analysis

Mouse lungs were fixed in formalin and embedded in parafilm blocks following standard method [24]. Once sliced, the lung section was deparaffined and rehydrated. Lung inflammation was evaluated on sections stained using the Carstair method (Electron Microscopy Sciences, Hatfield, PA, USA). Stained sections were digitalized using an AxioScan Z1 (Carl Zeiss, Oberkochen, Germany) at 10X magnitude. Each lung image was split into six sub-sections where inflammation in the perivascular, peribronchial, and parenchymal areas was scored. Scores of 1 to 5 were attributed for each area by sub-section and the mean score from the sub-sections presented.

### Histological Immunofluorescence

Lung sections were processed as described above. Following deparaffination and rehydration, antigen retrieval was performed on tissue following the protocol described in our previous work [25]. Then sections were permeabilized with TBS with 0.025% Triton X100 twice times 5 minutes and blocked with TBS containing 1% BSA and 5-10% goat serum (Multicell Wisent Inc.) for two hours. Tissues were labeled with 10 μg/mL of rat anti-GR1 clone RB6-8C5 (Thermo Fisher Scientific), 5 μg/mL of rat anti-CD4 clone 4SM95 (ThermoFisher Scientific), 1.14 μg/mL of rabbit anti-CD8a clone EPR20305 (Abcam Inc., Toronto, ON, Canada), 25 μg/mL of rabbit anti-SARS-CoV-2-N (Rockland Immunochemicals Inc., Limerick, PA, USA) in TBS with 1% BSA and 2.5-5% goat serum for 16 hours at 4°C. Primary antibodies were stained with eider 4 μg/mL of goat anti-rabbit-Alexa-488 (Thermo Fisher Scientific), 1.2 μg/mL of goat anti-rabbit-Alexa-647 (Thermo Fisher Scientific), or 0.7 μg/mL goat anti-rat-Alexa-647+ (Thermo Fisher Scientific) for 1 hour at room temperature in TBS with 1% BSA and 2.5-5% goat serum. Sections were incubated into TBS with 1.67 μg/mL of DAPI (Invivogen) then washed for 5 min in TBS and mounted with ProLong Glass Antifade reagent (Thermo Fisher Scientific). Immunostained sections were scanned using an AxioScan Z1 (Carl Zeiss) at 20X magnitude. Images were exported from Zen Blue (Carl Zeiss) to 8-bit TIFF. Stained cells were quantified using macro generated using Celeste Image Analysis Software (Thermo Fisher Scientific) and SARS-CoV-2 N positive areas were quantified using Fiji/ImageJ [26] and then normalized with the tissue area analyzed. The total tissue areas for normalization were measured using raw data czi file in Zen Blue using the DAPI channel.

### Cytokines, Chemokines, and Interferon Quantification

The protein inflammatory mediators were quantified using ProcartaPlexTM Mouse Mix & Match Panels kit (Thermo Fisher Scientific) on lung homogenate in the mice section.

### Lipidomic analysis

Mouse lungs were collected and homogenized as described in the mice section. Lung homogenates were heated at 65°C for 30 minutes to inactivate infectious viruses before being taken out of the BSL3 facility. Oxylipin extractions and analysis were carried out as described elsewhere [27]. Briefly, 10 μL of a deuterated standard was combined with 200 μL of lung homogenates prior to freeze-drying. A volume of 289 μL of an ethanolic solution (41%) was then added and mixed thoroughly to reconstitute the samples. The specimens were incubated with acetonitrile (2:1) at room temperature and then at -20°C in order to precipitate the proteins. After centrifugation, supernatants were removed and combined with 600 μL of an ammonium hydroxide solution (0.01 M) before being loaded on solid phase extraction columns. A portion of the unwanted compounds were washed from the cartridges with the aforementioned solution of ammonium hydroxide and then a mixture of acetonitrile and methanol (8:2). Compounds were eluted using an acidified version of the latter organic solution, nitrogen dried and reconstituted in 60uL (30% acetonitrile with 0.01% acetic acid). In order to enhance the precision of the quantification, a spiked calibration curve was generated by using 10 μL of each final sample to form a pool, which was employed to reconstitute pure standards extracted with the same steps as the samples. Analysis was performed with a High-performance liquid chromatography-tandem mass spectrometry system operated in negative mode and using specific transitions in scheduled multiple reaction monitoring (Table 1S) previously described [28]. The statistical analyses were performed using lipid concentrations normalized to total protein quantification. Then data were imported into MetaboAnalysis 6.0 online tools where they were log10 transformed and range scaled before any statistical analysis. The One-factor statistical analysis module was used to compare each mouse group for each time point using an unpaired t-test assuming an equal group variance. The metadata table statistical analysis module with covariates design was used to highlight the interactions between age or infection and expression of different lipids. A Two-way ANOVA was used to highlight the interaction between lipid expression and covariates (Age or infection time point). Linear models with adjustment for infection covariate were used to characterize the relation between age and lipid expression.

## Results

### COVID-19 course in K18-hACE2 elderly and young adult mice

As shown in figure 1A, mice with a mean age of 1.7 months (young group) or 18.5 months (old group) were infected intranasally with 500 TCID_50_ of SARS-CoV-2 and monitored for weight loss and overall appearance for 14 days. Mice that reached ethical checkpoint limits (>20% weight loss plus one other signs of discomfort) were euthanized. As shown in figure 1B, infected young mice experienced greater weight losses, especially on days 6-7 post-infection (DPI) relative to aged mice. However, while young mice stopped losing weight after day 7, older mice gradually continued losing weight, reaching endpoint limits. As a result, none of the aged mice survived the infection passed day 10 while 30% of young mice did (Figure 1C). However, likely due to the small number of animals used (n=10/group), no statistical differences in survival between the two groups were observed.

**Fig 1.**
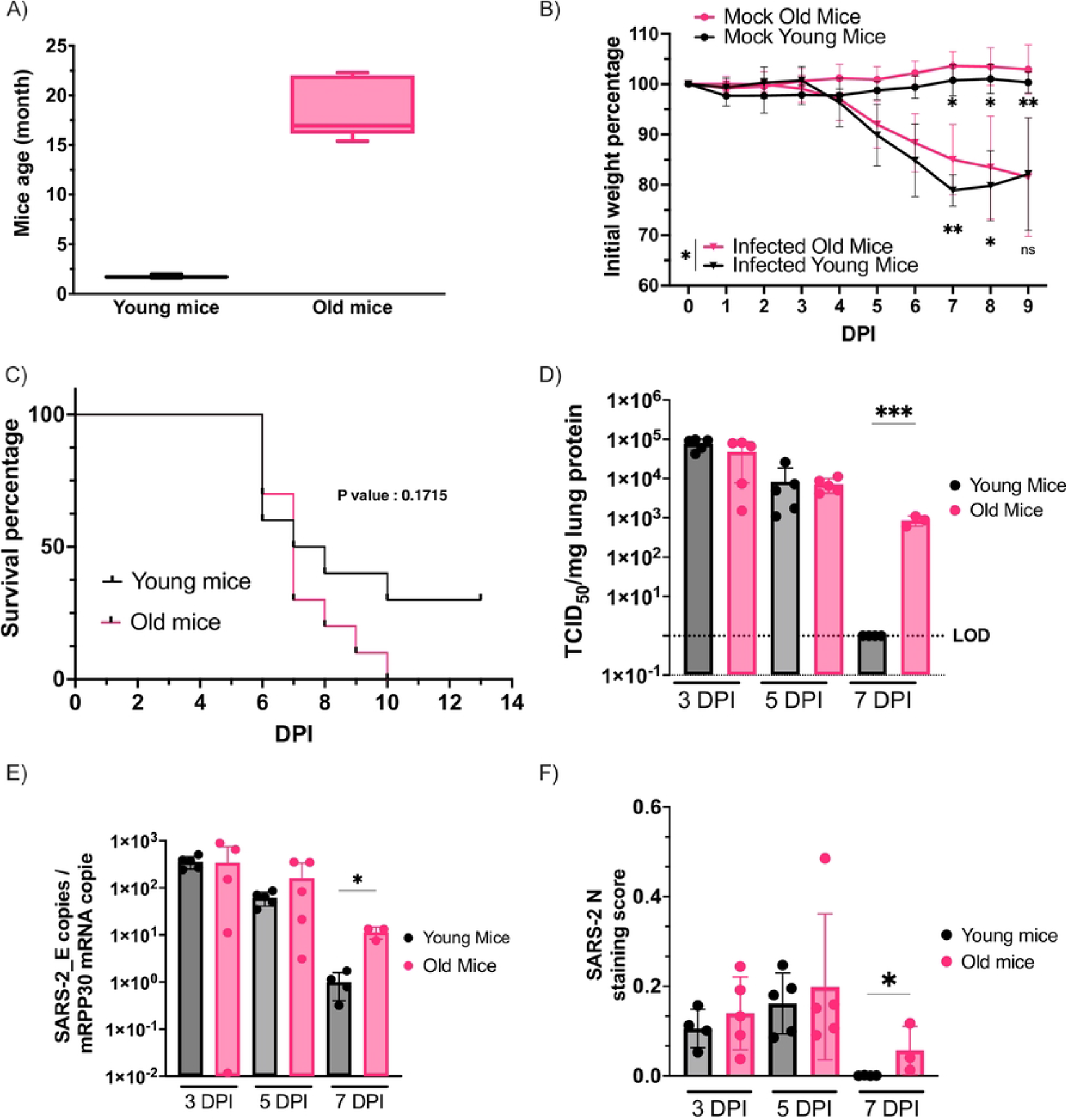
SARS-CoV-2 pathogenesis and viral replication in aged and younger K18-hACE2 mice. (A) Mice age distribution across young and old groups in months. Error bars display minimum and maximum age within groups, when bare in the boxplot represents the median age. (B) Mouse weight variation following mock as a function of time post infection. Values represent mean +/-SD expressed in percent relative to day 0 weight. The weight curves of both SARS-CoV-2 infected groups were compared using Two-way ANOVA. Uncorrected Fisher’s LSD tests were used to evaluate weight loss recovery by comparing weight at 7 to 9 DPI with the weight from 6 DPI that correspond to the weight loss peak for young mice. (C) Mice survival curve following infection with SARS-CoV-2. Mice were sacrificed when reaching the ethical checkpoints and those that reached the 14 DPI were considered as survivors. The difference in the survival was assay using a Log-rank (Mantel-Cox) test. (D) Lung homogenate infectious viral loads at 3 DPI to 7 DPI expressed in Tissue Culture Infectious Dose 50 by mg of lung protein (TCID_50/mg_) (Mean ± SD) and LOD line show the limit of detection of the assay. Comparison between groups was performed using an Unpaired T-test. (E) Viral RNA quantitation in the lungs using RT-ddPCR expressed in copies of SARS-CoV-2 E gene relative to copies of mRPP30 mRNA housekeeping gene (Mean ± SD). Comparison between groups was performed using an Unpaired T-test for 3 and 5 DPI or Mann Whitney test for 7 DPI. (F) SARS-CoV-2 N antigen quantification on lung section. Staining scores represent the proportion of tissue area with a positive signal (Mean ± SD). Comparison between groups was performed using an Unpaired T-test for 3 and 5 DPI or Mann Whitney test for 7 DPI. P value: <0.05 (*), <0.0021 (**), <0.0002(***), non-significant (ns).

### Deficient viral clearance in old mice lungs

Following virus inoculation, five mice per group were sacrificed on day 3, 5, and 7 post-infection and their lungs harvested for viral loads determination. Infectious virus titers are displayed in figure 1D. No significant viral load difference was detected between groups during the acute infection phase (3-5 DPI). However, a major difference was observed on day 7. While all young mice had viral loads that were below the detection limits, none of the elderly mice efficiently cleared the virus. Reduced viral clearance by aged mice was confirmed using viral RNA (Figure 1E) and N antigen (Figure 1F) measurements.

### Young mice produce chemokines in greater quantities at early times post infection relatively to aged mice

From the mouse lung homogenates obtained at 3, 5 and 7 DPI, several inflammatory mediators were measured. As shown in Figure 2A, the total inflammatory burden was significantly higher at 3-5 DPI in young mice relative to aged mice. On day 7, aged mice appear to produce overall higher levels of cytokines in response to infection. This result should be interpreted with caution as one of the three remaining mise over produced IL-6 (supplementary figure 1). When analyzed separately, CCL2 and CXCL9 were the main chemokines produced at greater levels by young mice at early time points (3 and 5 DPI) (figure 2B-E). In response to SARS-CoV-2 infection, CCL3 levels were produced at significantly higher levels in aged mice on day 7 post-infection relative to young mice (Figure 2F).

**Fig 2.**
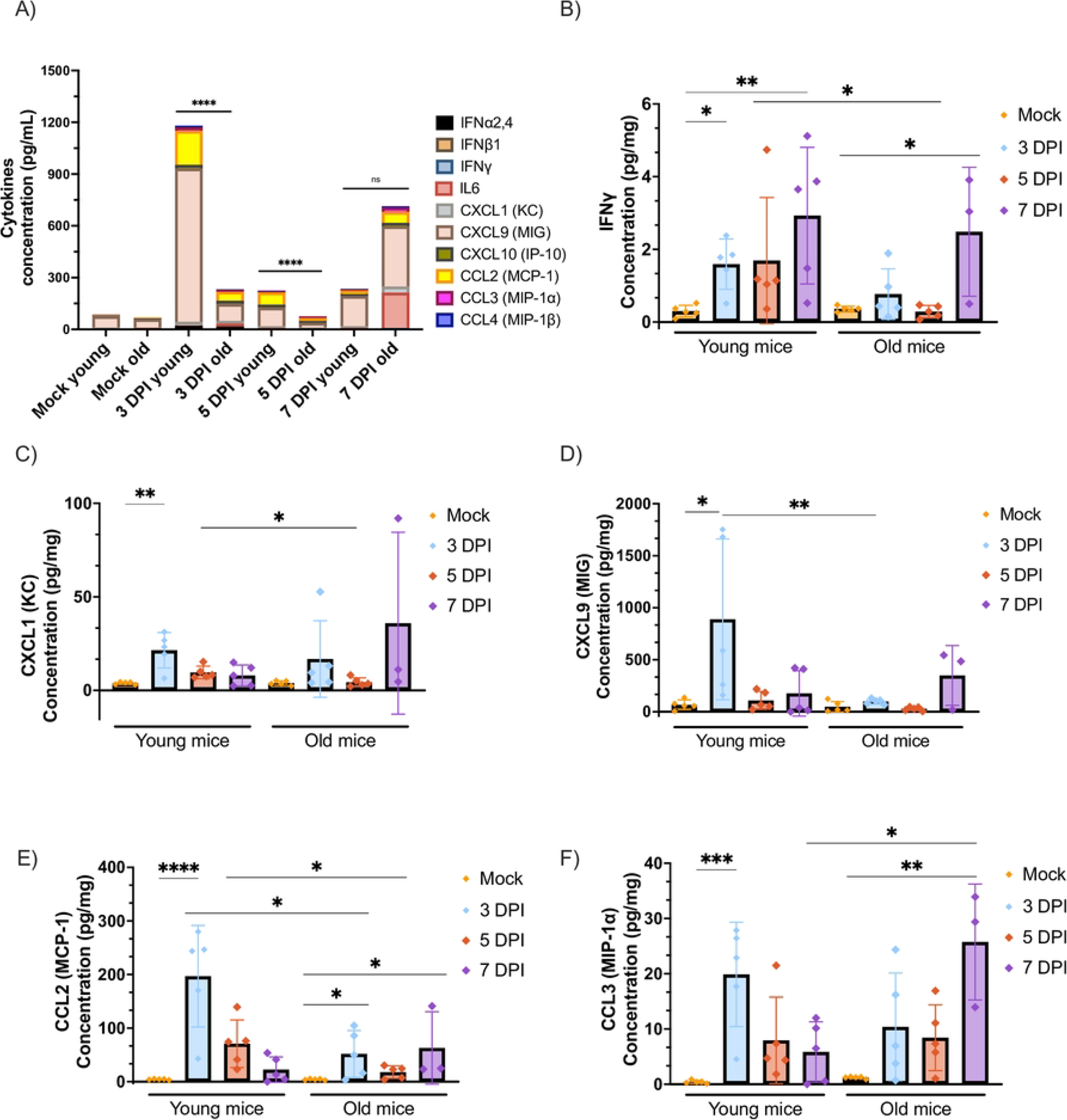
Chemokines, cytokines, and interferons (IFN) production in the lung following SARS-CoV-2 infection. Mediators were quantified from lung homogenates using Luminex technology (ProcartaPlex kit). (A) Total protein mediator burst. The subsections of each bar graph represent the mean concentration value for one mediator expressed in pg of mediator by mg of lung protein. Groups were compared using an Uncorrected Fisher’s LSD on the mean concentration by bare. IFNγ (B), CXCL1 (C), CXCL9 (D), CCL2(E), and CCL3 (F) production expressed in pg of mediator by mg of lung protein (Mean ± SD). For each group infection time points were compared with an Uncorrected Fisher’s LSD (young mice) or Uncorrected Kruskal-Wallis test (old mice) with their mock condition. Time points between groups were compared with the Unpaired T-test (3 and 5 DPI) or Mann Whitney (7 DPI). P value: <0.05 (*), <0.0021 (**), <0.0002(***) and <0.00001(****).

### No significant difference in the induction of proinflammatory cytokine and Type I-II IFN production or ISG expression

As for other cytokines/cytokines, IL-6 production was quantified in the lung homogenate of the different mouse groups, but no significative differences were observed (supplementary figure 1). No increase in IFN gene expression was noted (figure 3A). Similarly, CCL4, CXCL10 and IFNα(2,4) were highly induced by the infection without being significantly differently expressed between groups (Figure supplementary 1 A to E). *Il18, Ifnl2-3* and ISGs (IRF7, Isg56 and Isg15) genes expression was also induced by infection but did not differ between young and old mice (figure 3D-G). In contrast, *Tnf* gene expression was reduced in aged mice at 3DPI relative to young mice (Figure 3C).

**Fig 3.**
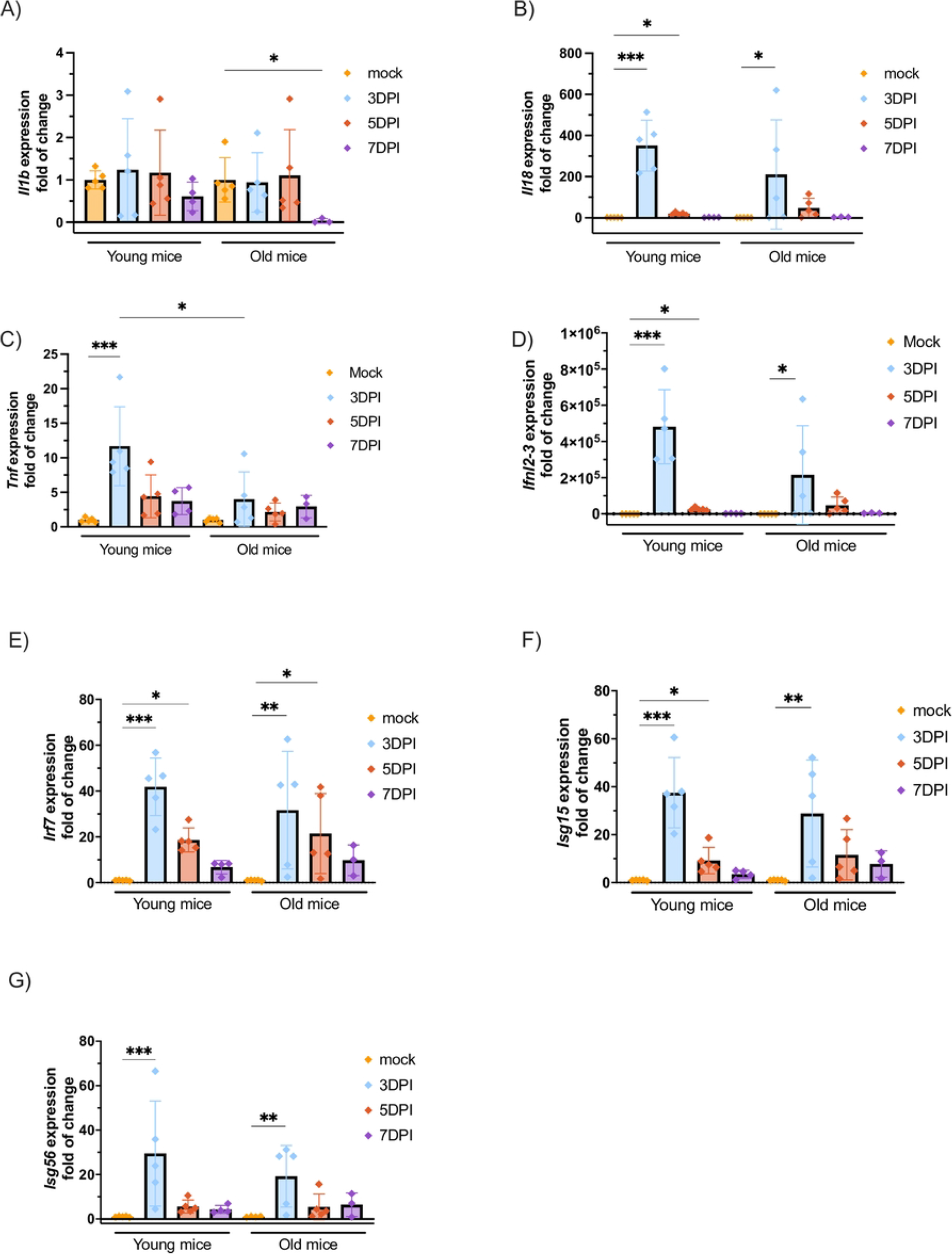
Inflammatory and antiviral genes expression levels in the lungs following SARS-CoV-2 infection. Gene expression was quantified using RT-qPCR from lung total RNA. *Il1b*(A), *Il18*(B) *Tnf*(C), *Ifnl2-3*(D), *Irf7*(E), *Isg15*(F), *Isg56*(G) gene expression were reported as fold of change in comparison with the respective mice group mock controls (Mean ± SD). For each group infection time points were compared with an Uncorrected Fisher’s LSD or Uncorrected Kruskal-Wallis test (old mice) with their mock condition. Time points between groups were compared with the Unpaired T-test (3 and 5 DPI) or Mann Whitney (7 DPI). P value: <0.05 (*), <0.0021 (**) and <0.0002(***).

### Elderly mice showed fewer reduced signs of inflammation at early time points after infection

Using the lung section stained using the Carstair method, the lung was evaluated for perivascular, peribronchial, and parenchymal signs of inflammation. As shown in Figure 4A, when the whole lung inflammation scores were tabulated, increased inflammation was observed in the lungs of young mice at 3 DPI and 5 DPI in comparison with aged mice. However, only peribranchial (3 DPI) and parenchymal (3 DPI and 5 DPI) zones showed a significantly lower inflammation level in old mice (Figure 4 B and C).

**Fig 4.**
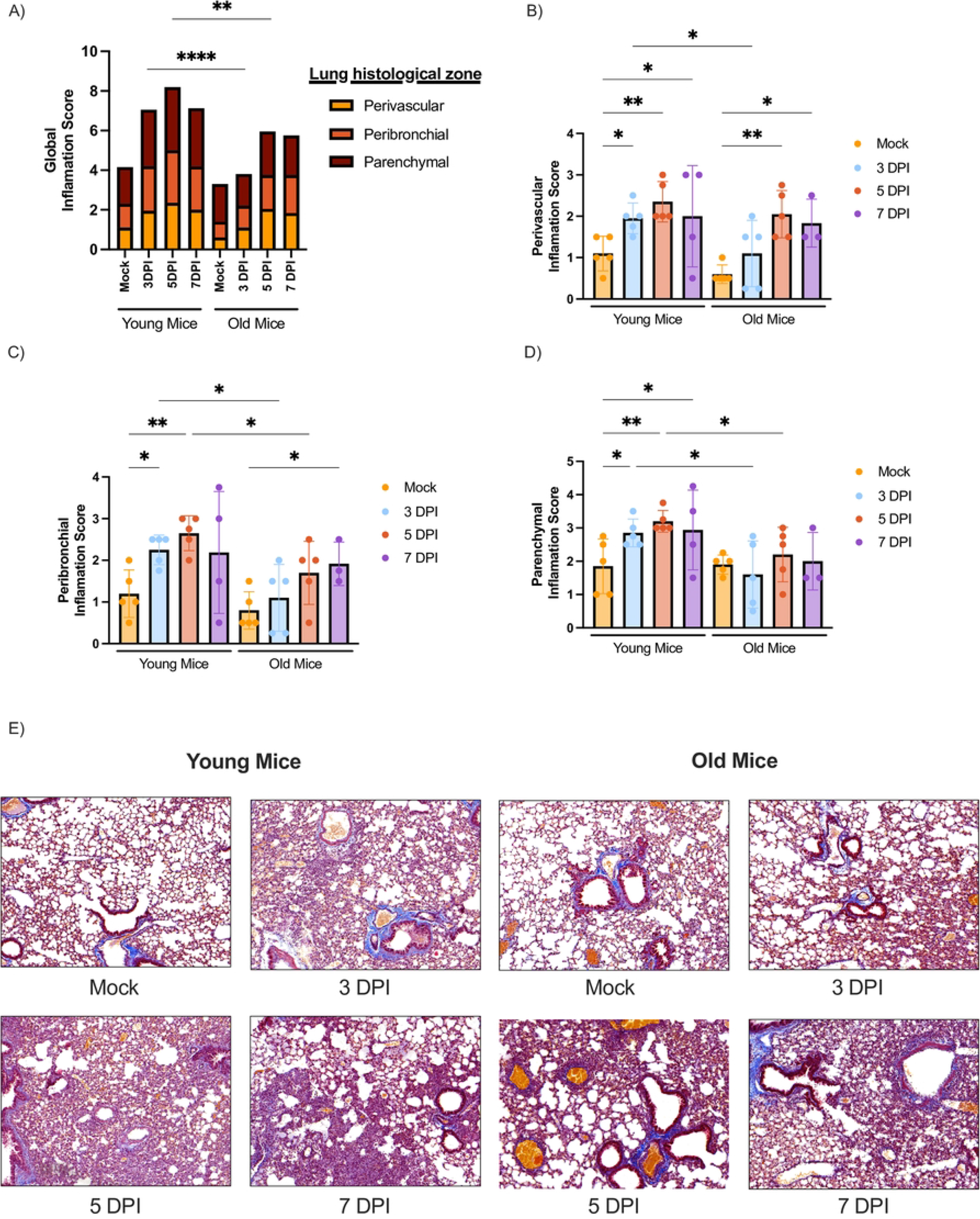
Lung histological inflammation in K18-mice in response against SARS-CoV-2 infection. Carstair stained sections were scanned and scored for inflammation (0 to 5) in the three different lung regions (perivascular, peribronchial, and parenchymal). Global inflammation score (A). Each subsection of the stacked bar graph represented the respective score for each lung region. Perivascular (B), peribronchial (C), and parenchymal (D) individual inflammation scores scaled from 0 to 5. Mouse groups and infection time points were compared using Uncorrected Fisher’s LSD. P value: <0.05 (*), <0.0021 (**), <0.0002(***) and <0.00001(****). Image representative of a Carstair stained lung section for each mice group was shown in (D).

### Younger mice exhibit a higher lung lymphocyte infiltration relatively to old mice in response to SARS-CoV-2 infection

Lung sections were also used to analyze the subpopulations of leucocytes following infection. CD4 marker was used to quantify T helper cells, CD8α was used to quantify cytotoxic T cells, and GR1 (Ly-6C/Ly-6G) was used as a marker of granulocytes. As shown in Figure 5A, the number of CD4 cells in mock infected animals was higher in young versus aged mice. Upon infection, a reduction in the number of CD4-positive cells in the lungs of younger mice but not in older mice was observed. Similar to CD4+ T cells, mock infected young mice had higher numbers of CD8+ T cells than aged mice, although not statistically different. In contrast, mock-infected aged mice had higher levels of resident CD8+ than young mice. Infection led to a significant increase in CD8+ T cells in young mice but not in old mice. (Figure 5B). Relative to young mice, the CD8 cell count was significantly lower at 5 DPI for aged mice. Regarding granulocytes, no recruitment was detected on 5 DPI for both groups (Figure 5C). Relative to mock infected mice, an increase of GR1 positive cell count was observed in aged mice at 7 DPI but not for younger mice.

**Fig 5.**
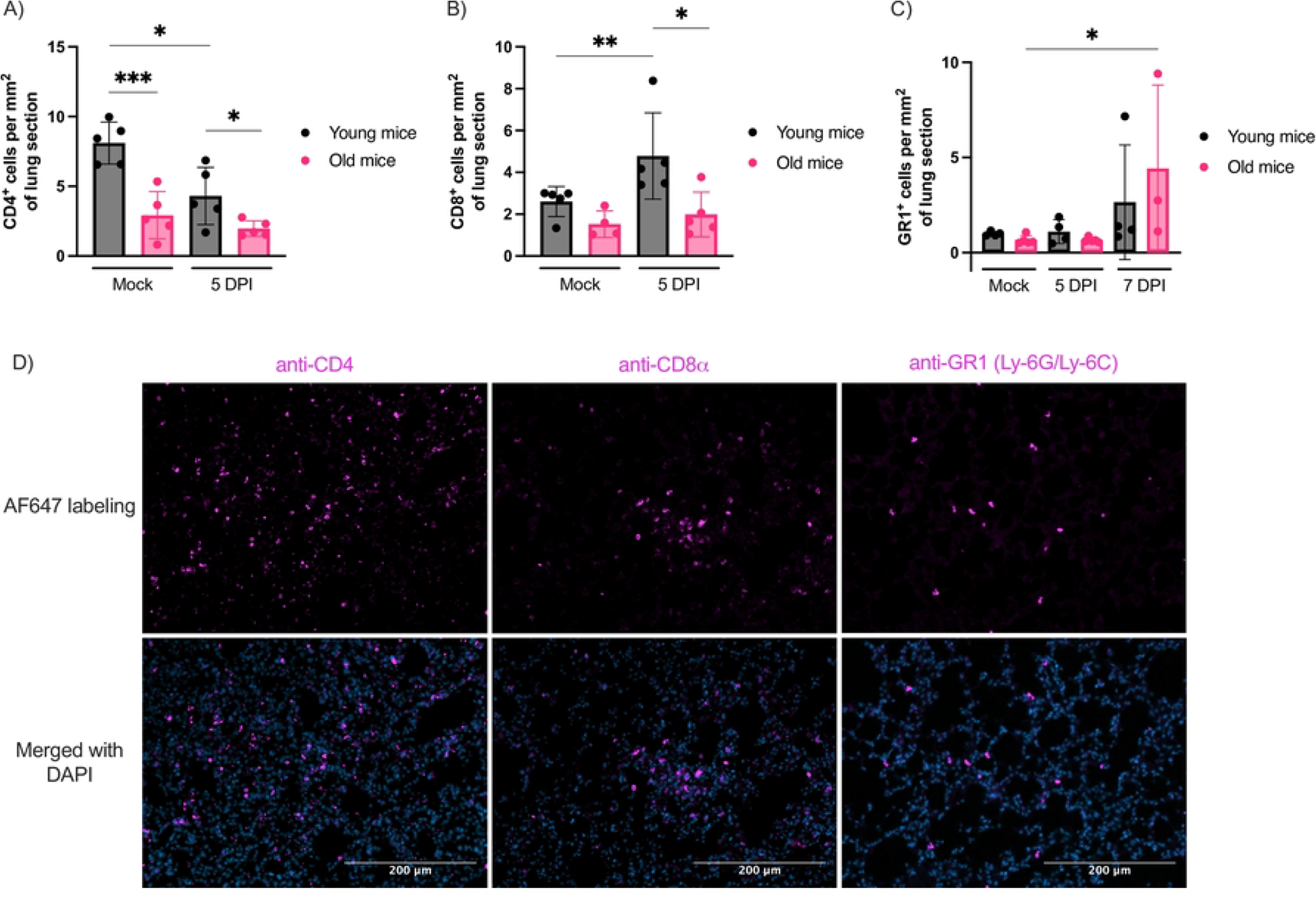
Lymphocytes and granulocytes recruitment in mouse lungs following SARS-CoV-2 infection. Lung sections were stained with antibodies against mice CD4 (A), CD8α (B), or GR1 (C). Stained tissues were scanned, and positive cells were quantified on the whole section scan. Results were expressed in a number of positive cells by mm^2^ of tissue section (Mean ± SD). Groups were compared with an Unpaired T-test or Mann Whitney according to the data distribution. P value: <0.05 (*), <0.0021 (**) and <0.0002(***). Images representative of each immunostaining was shown in (D).

### Aged mice showed an altered bioactive lipid basal profile and differential regulation than young mice in response to SARS-CoV-2 infection

We have extracted lipids from the mouse lung to measure the lipid mediators of inflammation (LMI) produced in response to infection. The lipidome was quantified using a panel containing a hundred lipids (listed in Supplementary Table 1). Those lipids are produced from arachidonic acid (AA), eicosapentaenoic acid (EPA), docosahexaenoic acid (DHA), linoleic Acid (LA) or α-linolenic acid (ALA) via cyclooxygenases (COX), lipoxygenase (LOX), cytochrome P450 (CYP) or non-enzymatic reactions (Figure 6). Lipids’ relative levels across mouse groups are presented in a bubble plot. Of quantified lipids, approximately 40 differed in elderly mice comparatively to the juveniles in at least one infection time-point (Figure 7A). When comparing the elderly and juvenile mock groups, ±11-/8-HEPE, ±8(9)DiHETE, 8(9)EpETE, 9(S)-HOTrE, prostaglandin E2 (PGE_2_), 8-iso-PGE2 and thromboxane B2 (TBX_2_) were detected at significantly higher levels in the elderly than young mice. On day 3 post-infection, ±11- or 15-HEPE, and ±14(15)-DiHETE derived from EPA were observed at significantly higher concentrations in older than young animals. Concomitantly LA/ALA derivatives like 13S-/9S-HOTrE or DIHOMEs and, DHA derivatives like resolvin D3 (RvD3), protectin Dx (PDx), ±16(17)-DiHDPA, 4-series neuroprostanes (4-F_4t_-NeuroP), 5-series isoprostanes (5-(*RS*)-5-F_2c_-IsoP) ±14 were also more prevalent in older than young animals. In opposition to other time points, at day 5 post-infection a cluster of decreased lipids concentration was observed in aged mice. This cluster contains lipids belonging to the HETEs, HEPEs, DiHDPAs, EpDPAs, HDPAs, and OxoETEs together with RvD2 and PGF_3□_ For the day 7 post-infection due to the small sample size and high variability, only 17(18)-DiHETE and 8(S)-HETE levels were differentially changed between the two mouse groups.

**Fig 6.**
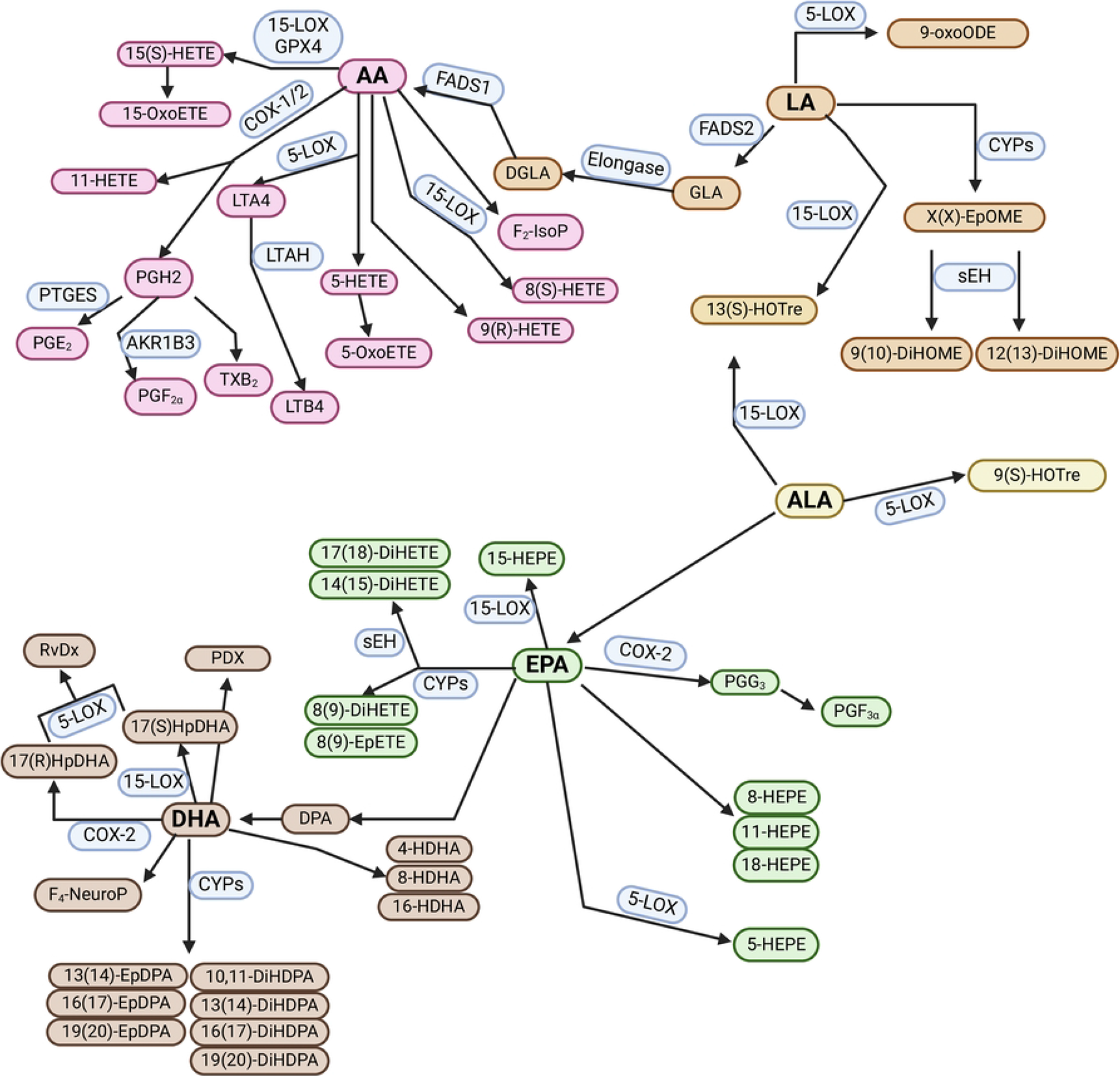
Production pathways of differentially expressed lipids (DELs) between elderly and young mice infected of not (mock) with SARS-CoV-2. These lipids are derived from Arachidonic Acid (AA), Eicosapentaenoic acid (EPA), Docosahexaenoic acid (DHA), Docosapentaenoic acid (DPA), Linoleic Acid (LA) alpha-Linolenic acid (ALA), gamma-Linolenic acid (GLA), Dihomo-gamma-Linolenic acid (DGLA) using enzyme Cyclooxygenases (COX), lipoxygenase (LOX), cytochrome P450 (CYP), Soluble epoxide hydrolase (sEH) or non-enzymatic reaction. Fatty Acid Desaturase (FADS), Elongase, Glutathione peroxidase 4 (GPX4), Leukotriene-A4 hydrolase (LTAH), Prostaglandin E synthase (PTGES), Aldose reductase (AKR1B3) enzymes were also needed to produce some of those metabolites from Leukotriene A4 (LTA4), Prostaglandin H (PGH), Prostaglandin G (PGG), Isoleukotoxin/Leukotoxin (EpOME), Hydroperoxy Docosahexaenoic Acid (HpDHA). DELs identified belong Hydroxy Eicosatetraenoic Acids (HETE), Oxo Eicosatetraenoic Acid (oxoETE/KETE), Prostaglandin E (PGE), Prostaglandin F (PGF), Thromboxane B2 (TXB_2_), Leukotriene B4 (LTB_4_), Isoprostane (IsoP), Hydroxy Octadecatrienoic Acid (HOTre), Isoleukotoxin/Leukotoxin diol (DiHOME), Hydroxy Eicosapentaenoic Acid (HEPE), Dihydroxy Eicosatetraenoic Acid (DiHETE), Epoxy Eicosatetraenoic Acid (EpETE), Hydroxy Docosahexaenoic Acid (HDPA/HDoHE), Epoxy Docosapentaenoic Acid (EpDPA/EpDPE), Dihydroxy Docosapentaenoic Acid (DiHDPA/DiHDPE), Neuroprostane (NeuroP), Resolvin D (RvD), Protectin DX (PDX) sub-class of lipids. Production pathways were built using the KEGG and WikiPathways (LIPID MAPS) database and work from Coras & al., Calder, Wen [28-31]. The figure was created with BioRender.com.

**Fig 7.**
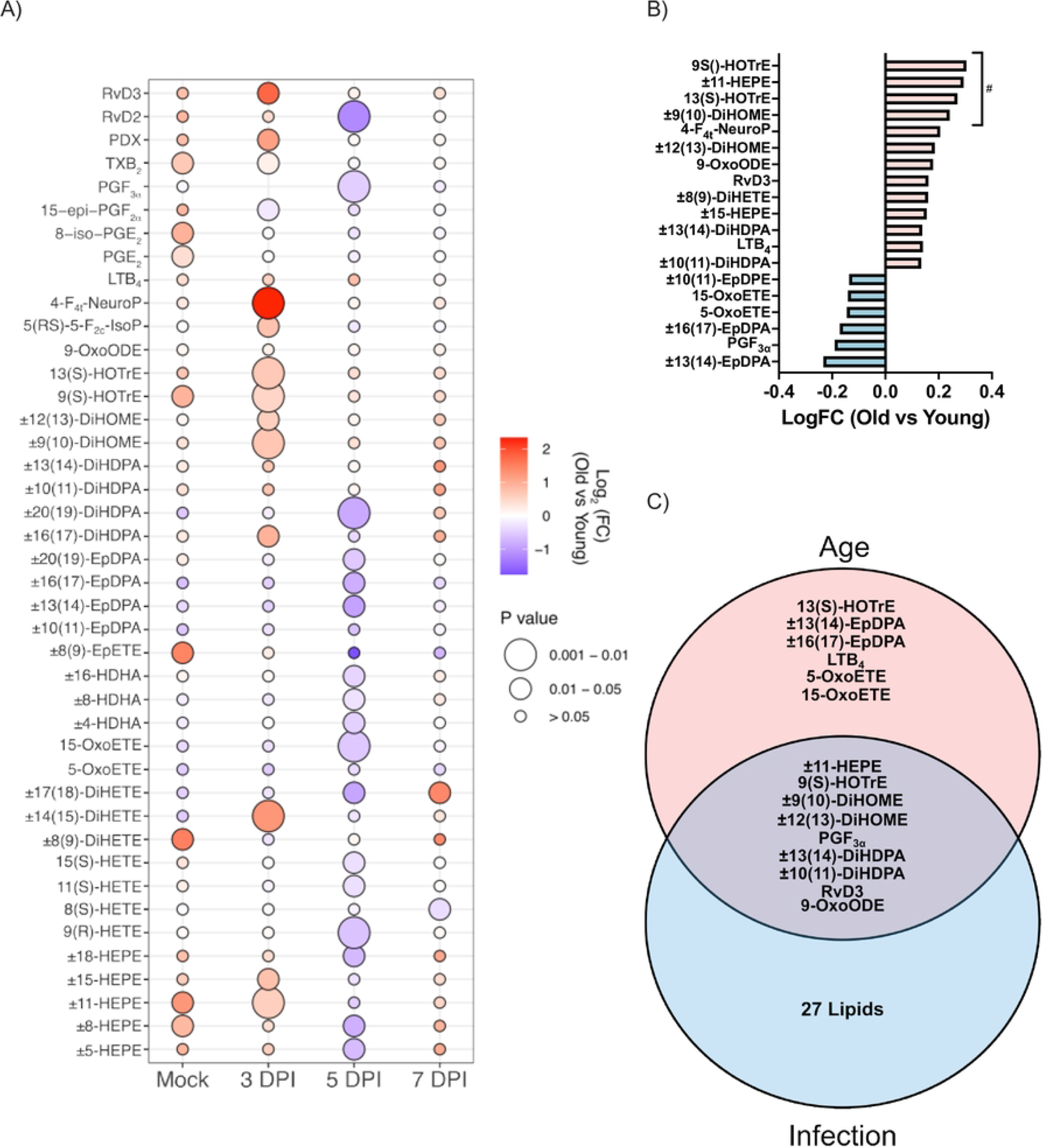
Lipidomic profiling of lung homogenates from the elderly and young mice infected or not (mock) with SARS-CoV-2. (A) Differentially expressed lipids (DELs) between the two mouse groups for the same infection time-point (mock, 3 DPI, 5 DPI, 7 DPI). Color blue to red indicates log_2_ of the fold change in the aged mice in comparison with the young mice and the dot size indicates the P value calculated with uncorrected unpaired T-test assuming equal group variance. (B) Levels of Lipids significantly different according to age when adjusted (marked with #) or not with the infection time-point. Lipids with a unadjusted P value below 0.05 in the bar graph. P values were calculated with a linear model based on the limma method [32]. (C) Specific Lipids showed a significant interaction with age and infection time-point. Lipids that display a raw P value below 0.05 at the Two-way ANOVA tests are shown in the Venn diagram. The 27 lipids that show no only interaction and differed only with the stage of infection were ±19(20)-DiHDPA, ±8-HEPE, ±16-HDPA, ±17(18)-DiHETE, ±20-HDPA, Prostaglandin E3, RvD2 n-3 DPA, ±5-HEPE, ±7(8)-DiHDPA, ±13-HDPA, 13(S)-HODE, ±8-HDHA, 9(S)-HODE, 9(R)-HETE, ±14(15)-DiHETE, 11(S)-HETE, ±5(6)-DiHETE, Thromboxane B2, ±(18)-HEPE, 17(S)-HDHA, ±4-HDPA, ±7-HDPA, ±11-HDPA, Protectin DX, 8-isoPG E2 and 5-F_2t_-IsoP.

Following a linear model analysis, our results show that 19 lipid mediators display a significant difference with the advanced age (old vs young) when adjusted or not for infection time point (Figure 7B). Among these lipids, 9(S)/13(S)-HOTrE, ±11-HEPE, and ±9(10)-DiHOME have shown significant difference only when adjusted with the covariate infection time point. Six lipids globally reduced with old age, and they were produced from DHA or EPA via CYP or AA using LOX15/5. The 13 other mediators showed a significant positive association, and they were mainly produced from EPA, DHA, AA, or LA/ALA through the actions of CYP, LOX, or via non-enzymatic reaction. When analyzed in an ANOVA 2 for interaction between age (old/young) and infection time point, six mediators were mainly influenced by age (Figure 7C). Most of them displayed were negatively impacted except 13(S)-HOTrE and LTB_4_ which showed a significant positive association (Figure 7B). The level of nine lipids was significantly impacted by both, infection and age and 27 lipids only by the infection. A summary of the differential lipid metabolite functions is shown in Supplementary Table 2.

## Discussion

Following SARS-CoV-2 infection, both young and aged K18 mice demonstrated signs of infection including ruffled hair and weight losses. However, elderly mice failed to recover their weight by 9 DPI and experienced a tendency for a worse survival rate. Differences in the severity of COVID-19 are less age-dependent in the K18-hACE2 model than what was observed in human patients [21]. This discordance between this model and humans is likely attributable, at least partially, to the wider tissue distribution and overexpression of the hACE2 receptor in K18 mice that lead to very high susceptibility of these mice to SARS-CoV-2 [33].

While young mice successfully cleared the infection by day 7, significant infectious viral loads were present in the lungs of aged mice. A similar viral persistence in elderly human patients was suggested previously by Longtin and collaborators [34]. Considering viral loads and type I and III IFN responses were similar in young and aged mice at early time points (days 3 and 5), this suggests that the viral persistence results from improper viral clearance rather than increased viral replication in aged mice.

When CCL2, CCL3, CXCL9, and IFNγ were measured in lung homogenates, older mice produced less such mediators in response to infection relative to young mice. Those mediators are well known as recruiters and activators of several leucocyte populations involved in the T cell response build-up [35-37]. More precisely CXCL9 and IFNγ are associated with a T helper (Th) 1 response that favors the elimination of intracellular pathogen-like virus [38]. Despite not being directly involved in the Th1 polarisation, CCL2 and CCL3 are associated with responses against viral infections and could favor the build-up of a potent antiviral response through the recruitment of NK cells and monocytes/macrophages [39]. *Tnf* gene expression was also reduced in infected old mice supporting the hypothesis of an impairment of the Th1 response in this group. Weaker production of those mediators could explain the viral persistence observed at 7 DPI in elderly mice. A reduced level of CXCL1 was also measured, but it did not impact negatively the granulocyte recruitment in the lung, which was known as the main target of this chemokine [40, 41].

As highlighted above, the global level of protein mediators (Cytokine, Chemokine, IFN) was higher in younger mice at 3-5 DPI relative to older mice. This finding correlates with higher global inflammatory scores for the same time points in the young mouse group. For this group, inflammation scores increased with the same magnitude from mock to 5 DPI in the three lung compartments (blood vessels, bronchus, and parenchyma). For the same time points in the aged mouse group, a significant increase in inflammatory scores was observed only in the blood vessels region at 5 DPI. This suggests that in elderly mice, leucocytes recruitment was reduced and failed to efficiently migrate through the blood vessels to reach the infection site. This finding is also supported by the global immunosenescent observed in lungs from aged patients, where inaccurate migration and impaired anti-microbial function of lung leucocytes were observed [42].

As was observed in peripheral blood of severe COVID-19 patients, CD4^+^ cell count in mouse lungs were decreased during SARS-CoV-2 infection [43]. In our model, the SARS-CoV-2 infection increases the CD8^+^ cell count in K18-mice lungs contrary to what was reported in peripheral blood from humans. A decrease in CD4^+^ cells in aged mock mice was observed which is in accordance with T cell age-dependent depletion in humans [42]. The lower CD4^+^ and CD8^+^ at 5 DPI in elderly mice is consequent to the lower production of CCL2, CCL3, CXCL9, and IFNγ and could explain the impaired viral clearance in this mouse group.

In the absence of infection, elderly mice have higher levels of PGE_2_ and TXB_2_, two lipids associated with pro-inflammatory activities [44, 45]. LTB_4,_ also a pro-inflammatory lipid mediator, is expressed at higher levels in aged mice. In contrast, 15-, identified as an NF-κB response inhibitor, is expressed at reduced levels in aged mice [46, 47]. During infection, some HOTrEs and 11-HEPE were globally present at higher levels in the elderly than in young mice 3 days post-infection. These mediators are mainly connected with anti-inflammatory and pro-resolution responses [48-51]. Together, these results suggest that aged mice exhibit a higher basal level of inflammatory lipid mediators in comparison to young mice. This finding is in accordance with the high expression of stress-related genes observed in elderly human subjects [52].

Following SARS-CoV-2 infection, elderly mice also displayed a different lung lipidome than younger mice. Notably, 12(13)-DiHOME and 9(10)-DiHOME, also known as Leukotoxin diol, showed a positive relation with age and were at present at significantly higher levels in aged mice day 3 post-infection. As these mediators are closely related to lung inflammation and potential contributors to ARDS, our results suggest a possible explanation for the worse outcome observed in elderly mice [53-58]. Elevated levels of DiHOMEs in plasma have been linked with the severity of COVID-19 [59]. In contrast, a major part of upregulated and differentially expressed lipids (DELs) at 3 DPI in elderly mice are known to have protective or pro-resolution effects, such as RvD3, PDx, 4-F_4t_-NeuroP, HOTrEs, ±14(15)-DiHETE, ±15-HEPE, and ±11-HEPE [48-51, 60-66]. This protective inflammatory response could mitigate the inflammation burst linked with DiHOMEs production. In addition to the potent inhibitory effect of ±14(15)-DiHETE on NK cells, these mediators could contribute to the impairment of viral clearance [67]. In accordance with the high production of immunomodulator lipids in aged mice at 3 DPI, most of the DELs at 5 DPI were lower in old than young mice. Not surprisingly, part of them have moderate pro-inflammatory roles like PGF_3□_ [68], 11(*S*)-HETE, and 8(*S*)-HETE [69, 70]. However, most of them have also immunomodulatory functions, notably, RvD2, 19(20)-DiHDPA, HDPAs, 15-OxoETE, ±17(18)-DiHETE, and ±18-HEPE [60, 65, 66, 71-76]. Notably, the level of 15(*S*)-HETE was lower in 5 DPI aged mice than in their younger counterparts. The urinary 15-HETE expression in mild COVID-19 patients has been correlated with higher CD4 and CD8 T cell activation [59].

The differential regulation of anti-inflammatory mediators at 3 DPI and 5 DPI may suggest that the pro-resolution response takes a longer time to build up in younger mice allowing the establishment of a more efficient antiviral response and better infection management. Specific isomers of 5-series F_2_-isoPs and 4-series F_4_-NeuroPs, markers of oxidative stress, were much higher in old than younger mice 3 days post-infection as reported here in the lung. This is in accordance with plasma levels of 5-series and 14-series F_2_-isoPs that were previously associated with the severity of COVID-19 infection in human patients [59, 77]. Increase in 5-series and 15-series F_2_-isoPs in plasma had also been associated with COVID-19 ICU admission[78]. Moreover, patients who survived or died from COVID-19 showed higher levels of 4-hydroxy-nonenal (4-HNE) a marker of lipid peroxidation in plasma than healthy individuals [79]. Our results are in line with the literature for increased oxidative stress in more severe infections found in old animals.

The difference in the lipidome profiles between aged and young mice did not match with the lipid mediator increased in hospitalized COVID-19 patient BALF or exhaled breath condensates. Indeed, increased levels of ±19(20)-EpDPA, 15(S)-HETE, ±15-HEPE, and ±5-HEPE in COVID-19 patients have been reported while our results suggest a lower level of these lipids in the lungs of older mice [15, 80]. These results confirm the difficulty of comparing mouse lung homogenates with COVID-19 patients BALF, as previously stated [25]. Severe COVID-19 in human patients tends to progress more slowly than in acute infection animal models such as K18-hACE2 mice. The median time from onset to hospitalization was 7 to 11 days and admission to an intensive care unit was 9 days [81-83]. In our previous work, a higher level of LTB4, PDX, and 13-HOTrE was detected in BALF from intubated COVID-19 patients which is in line with what was observed in aged mice. On the other hand, 5/15-HETE/OXOETE and RvD2 were detected at lower levels in elderly mice when highly produced in human patients [15]. Lower level of 5/15-HETE in the plasma had been associated with COVID-19 ICU admission which was in accordance with the decreased level of those compounds in aged mice [78]. Moreover, we detected an increased level of RvD3 and PGF_3□_ that was not observed in the Severe/ICU patient BALF or plasma[15, 78]. In our experiment setup, the mice began to show severe respiratory disease signs from day 5 post-infection, it is predictable that the lipidomic profile didn’t reflect what has been observed in hospitalized COVID-19 patients. Work from Wong & al. highlights the implication of the PGD_2_ pathways on the viral clearance and disease outcome in middle-aged mice infected with adapted SARS-CoV-2 strain [84]. Our work did not detect a change in the PGD_2_ expression in elderly infected mice in comparison with young ones. This may be explained by the different disease courses observed in wild-type C57BL/6 used in Wong’s studies versus the K18-hACE2 mice used in the present studies. As the K18 model is more susceptible to SARS-CoV-2, we observed a decrease in survival in young mice which is not the case with young wild-type C57BL/6 [84].

## Conclusion

In summary, the present study highlights that SARS-CoV-2 infection of K18-hACE2 mice triggers a robust immune response in younger mice characterized by a production of Th1-related mediators and antiviral chemokines. This response was significantly reduced in aged mice. Consequently, more severe clinical symptoms and deficient viral elimination were observed in the aged mouse cohort. Moreover, we observed a deep modulation of the bioactive lipid profile between the two mouse groups following SARS-CoV-2 infection. Indeed, lipids that were involved in anti-inflammatory and protective activity were detected at a higher level in aged mouse lungs at an early infection time-point. This finding suggests that the balance between immune activator and repressor signals given plays a crucial role in the build-up of an efficient antiviral response against SARS-CoV-2. Those results highlight the potential of targeting the immune response instead of the viral replication as a treatment against severe COVID-19 experienced by elderly people.

## Supporting information

**S1 Fig. Chemokines, cytokines, and Interferons (IFN) production following SARS-CoV-2 infection panel completion.** IL6 (A), CCL2 (B), CXCL10(C), IFNα2,4 (D), and IFNβ (D) production expressed in pg of mediator by mg of lung protein (Mean ± SD). For each group infection time points were compared with an Uncorrected Fisher’s LSD (young mice) or Uncorrected Kruskal-Wallis test (old mice) with their mock condition. Time-point between groups were compared with Unpaired T-test (3 and 5 DPI) or Mann Whitney (7 DPI). P value: <0.05 (*), <0.0021 (**), <0.0002(***) and <0.00001(****).

**S1 Table. Selected reaction monitoring parameters for tandem mass spectrometry optimized for each oxylipin.** DP: declustering potential, EP: entrance potential, CEP: collision cell entrance potential, CE: collision energy, CXP: collision cell exit potential, RT: retention time.

**S2 Table. Function and implication in disease of the lipids differentially produced.**

## Acknowledgments

Fluorescence and brightfield images of mice lung section were acquired using the slide scanner provided by the plateforme d’analyse d’images à haut-débit du centre de recherche du CHU de Québec-Université Laval.

## Ethics Statement

The study was conducted in accordance with the Declaration of Helsinki, and Mice protocols were approved by the Comité de protection des animaux de l’Université Laval. (Approval code: 22-1072).

